# Untargeted LC-MS metabolomics for the analysis of micro-scaled extracellular metabolites from *hepatocytes*

**DOI:** 10.1101/2020.09.24.312520

**Authors:** Rodi Abdalkader, Romanas Chaleckis, Isabel Meister, Pei Zhang, Craig Wheelock, Ken-ichrio Kamei

## Abstract

Metabolome analysis in micro physiological models is a challenge due to the low volume of cell culture medium (CCM). Here, we report a LC-MS-based untargeted metabolomics protocol for the detection of hepatocyte extracellular metabolites from micro-scale samples of CCM. Using a single LC-MS method we have detected 57 metabolites of which 27 showed >2-fold shifts after 72-hours incubation. We demonstrate that micro-scale CCM samples can be used for modelling micro-physiological temporal dynamics in metabolite intensities.

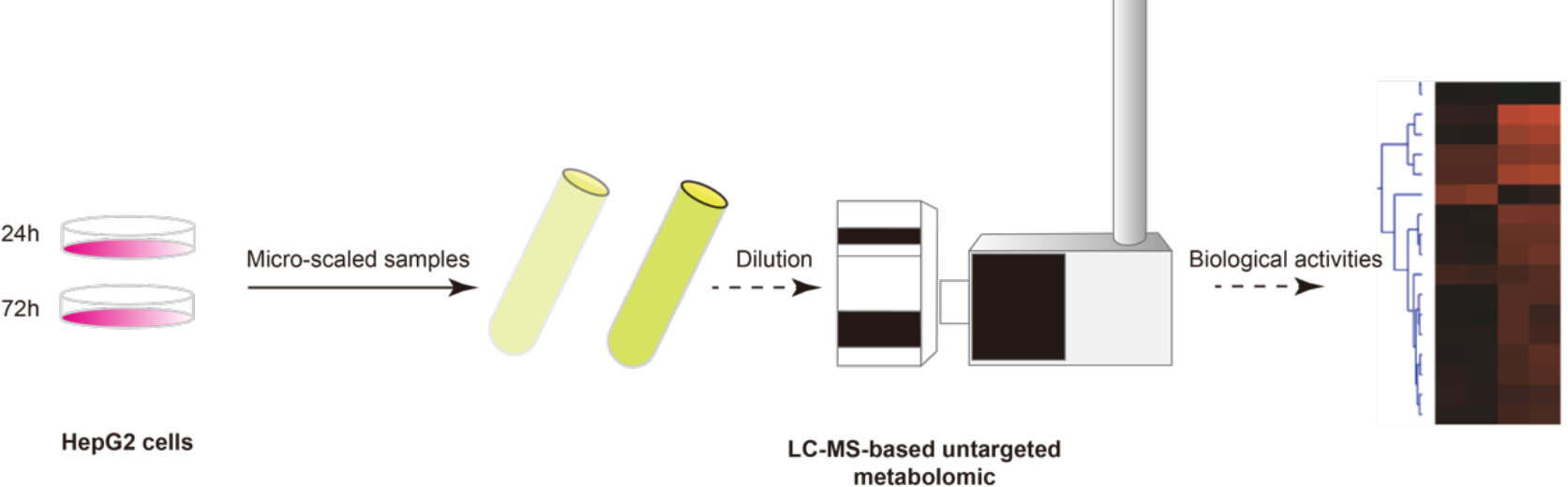
Graphical Abstract.

## Introduction

The liver is an essential organ for metabolism in the human body. Hepatocytes are the principal cells for metabolizing endogenous and exogenous compounds^1^. Also, they are the gold-standard model for addressing the absorption, distribution, metabolism, and excretion (ADME) of drugs ^2,3^. Recently there has been massive progress in the establishment of the micro-physiological models of the human liver in microfluidic devices that can replicate the anatomical structure of the liver as well as its blood flow ^4–7^. Moreover, the generation of human pluripotent stem cells (hPSCs)-derived liver mini-organoids is another emerging trend that has the potential to replace the classical two-dimensional hepatocytes models ^8,9^. Although, the microscale size of the liver on a chip or the liver organoid is considered an advantage for the high-throughput drug screening applications, yet it is also a challenge. The culturing volume inside the microfluidic channels is of only a few microliters per sample ^10^. Conventional analysis tools for the determination of metabolic activities such as the commercially available colorimetric and bioluminescent assays require much higher sample volume for the initiation of reaction between substrate and target metabolite. Furthermore, usually a single metabolite is detected per assay.

LC-MS-based untargeted metabolomics can detect a vast array of metabolites in a single measurement with relatively simple – yet efficient – sample preparation compared to other methods^11^. Advances in metabolomics now enable single-cell measurements^12^, however for cell culture medium (CCM), volumes of hundreds of microliters are still used for samples preparation^13^. There is no established methodology assessing the power of LC-MS-based untargeted metabolomics for detecting biological extracellular metabolites from a low micro-scaled volume of *in vitro* extracellular samples. Thus, the development of such LC-MS-based untargeted metabolomics platform will help us to understand detailed metabolic activities and their corresponding biological pathways at the microscale level of *in vitro* cell models. Here, we have investigated the capacity of LC-MS-based untargeted metabolomics for delivering temporal metabolite determination in micro-scaled volumes of extracellular metabolites secreted from the human hepatic cell line (HepG2). One microliter of CCM at 0, 24 and 72 h of cell culturing was extracted using 50 to 200 µL of acetonitrile. Only 1 µL of CCM was needed for the detection of 57 metabolites. Moreover, the biological impact of metabolite shifts was evaluated in relationship with the functions of the HepG2 cell functions.

## Experimental

### Cell culture and extracellular metabolite collection

HepG2 cells (American Type Culture Collection, Manassas, VA, USA) were cultured in DMEM supplemented with 10% (v/v) FBS, 1% penicillin/streptomycin, and 1 mM nonessential amino acids. The cells were passaged with trypsin-EDTA solutions at a 1:10 to 1:20 subculture ratio. For the collection of extracellular metabolites 3×10^5^ cells were seeded into T-25 flasks in a final volume of 15 mL of the CCM. Cells were then incubated at 37 °C and 5% CO_2_. At three timepoints (0 - control, 24 and 72 h) 1 mL of extracellular CCM was taken and preserved at −80 °C.

### CCM sample preparation, LC-MS metabolomics data acquisition and analysis

CCM samples were thawed at room temperature and 1 μL of each sample was extracted using 50, 100 μL or 200 μL acetonitrile containing five technical internal standards (tIS), Table S1. Then samples were centrifuged at 4°C for 15 min at 20000 g. Forty microliter of the supernatant were transferred to a 96-well 0.2 mL PCR plate PCR-96-MJ (BMBio,Tokyo, Japan). The plate was sealed with a pierceable seal (4titude, Wotton, UK) for 3 s at 180°C using a plate sealer (BioRad PX-1, CA, USA) and kept at 4°C during the LC-MS measurement. Injection volume was 1 μL (injection sequence Table S2). The LC-MS method is described in previous publications ^14–16^. Briefly, metabolite separation was performed on an Agilent 1290 Infinity II system using SeQuant ZIC-HILIC (Merck, Darmstadt, Germany) column using a 12 min gradient of acidified acetonitrile and water. Data was acquired on an Agilent 6550 Q-TOF-MS system with a mass range of 40−1200 m/z in positive ionization all ion fragmentation mode (AIF) including three sequential experiments at alternating collision energies: one full scan at 0 eV, followed by one MS/MS scan at 10 eV, and then followed by one MS/MS scan at 30 eV. The data acquisition rate was 6 scans/s. Data were processed in MS-DIAL version 4.20 ^17^ (detailed parameters in Tables S3 and S4). An in-house MS2 spectral library containing experimental MS2 spectra and retention times (RT) for 391 compounds obtained from standards^14,16^ was used for annotation of detected features using three criteria: (i) accurate mass (AM) match (tolerance: 0.01 Da), (ii) RT match (tolerance: 1 min), and (iii) MS2 spectrum match (similarity >70%). The MS2 similarity was scored by the simple dot product without any weighting (at least two MS2 peaks match with the reference spectra). The MS2 similarities with reference spectra were matched to any of the CorrDec^18^ or the MS2Dec^17^ deconvoluted MS2 spectra of the three collision energies (0, 10, and 30 eV). The dataset has been deposited to the EMBL-EBI MetaboLights repository with the identifier MTBLS1794 ^19^.

### Data visualization and statistical analysis

Data are represented as triplicate preparations and LC-MS measurements of samples. The unpaired multiple t test was performed using GraphPad prism 8 (GraphPad Software, La Jolla California, USA). Orange 3 software (Version 3.23.1; Bioinformatics Laboratory, Faculty of Computer and Information Science, University of Ljubljana, Slovenia^20^) was used for data mining. The biological pathways analysis was performed using the MetaboAnalyst platform^21^.

## Results and Discussion

For the development of micro-physiological models of the liver, the determination of the in situ metabolic activity is of critical importance. The measurement of extracellular metabolites in CCM enables longitudinal monitoring of the same cell culture. However, a huge obstacle in these models is the low micro-scaled volume in the cell culturing chambers that limits the number of detected metabolites. In this study, by using a single LC-MS method we have confidently identified 57 metabolites (Table S5) corresponding to the Metabolomics Standard Initiative MSI annotation level 1^22^: 32 were identified using accurate mass, retention time and MS2 spectral match (AMRT+MS2) and 25 were AMRT-matched. Four of the five tIS showed the coefficient of variation (CV) of peak areas <15 % across all CCM samples (Table S5). Median CV of all detected metabolites in the triplicate measurements was 8% (Table S5).

The analysis showed distinct trajectories of metabolite abundances in CCM during 72 h of the samples extracted using 100 μL acetonitrile (Fig. 1A). Eight metabolites showed >2-fold shifts between the 0 and 24 h samples and 27 metabolites showed >2-fold shifts between the 0 and 72 h samples (Fig. 1B). Comparable shifts were also observed in the samples extracted using 50 and 200 μL acetonitrile (Table S5). Extraction using 100 μL acetonitrile makes two measurement aliquots (40 μL) possible and provides reasonable signal intensities. Amino acids decreased over time indicating consumption, while modified purines and pyrimidines increased reflecting DNA metabolism. Biological pathways assigned by MetaboAnalyst related to these compounds confirm these observations by highlighting the significant involvement of aminoacyl-tRNA biosynthesis pathways, valine, leucine, isoleucine biosynthesis-degradation pathway, and purine metabolism pathway (Fig. 1C). These pathways are essential in hepatocytes for maintaining their characteristics as well as the metabolic activities within. For instance, amino acids are essential elements for the TCA cycle (tricarboxylic acid cycle) as energy generator in aerobic organisms^23^. These results indicate that of LC-MS-based untargeted metabolomics can be used for micro-scaled extracellular samples such as in organ-on-the-chip applications where only small sample volumes are available. We therefore envision the application of our methods for the in-situ analysis of the metabolic activities in micro-physiological models of the human organs in microfluidic devices.

**Figure 1.**
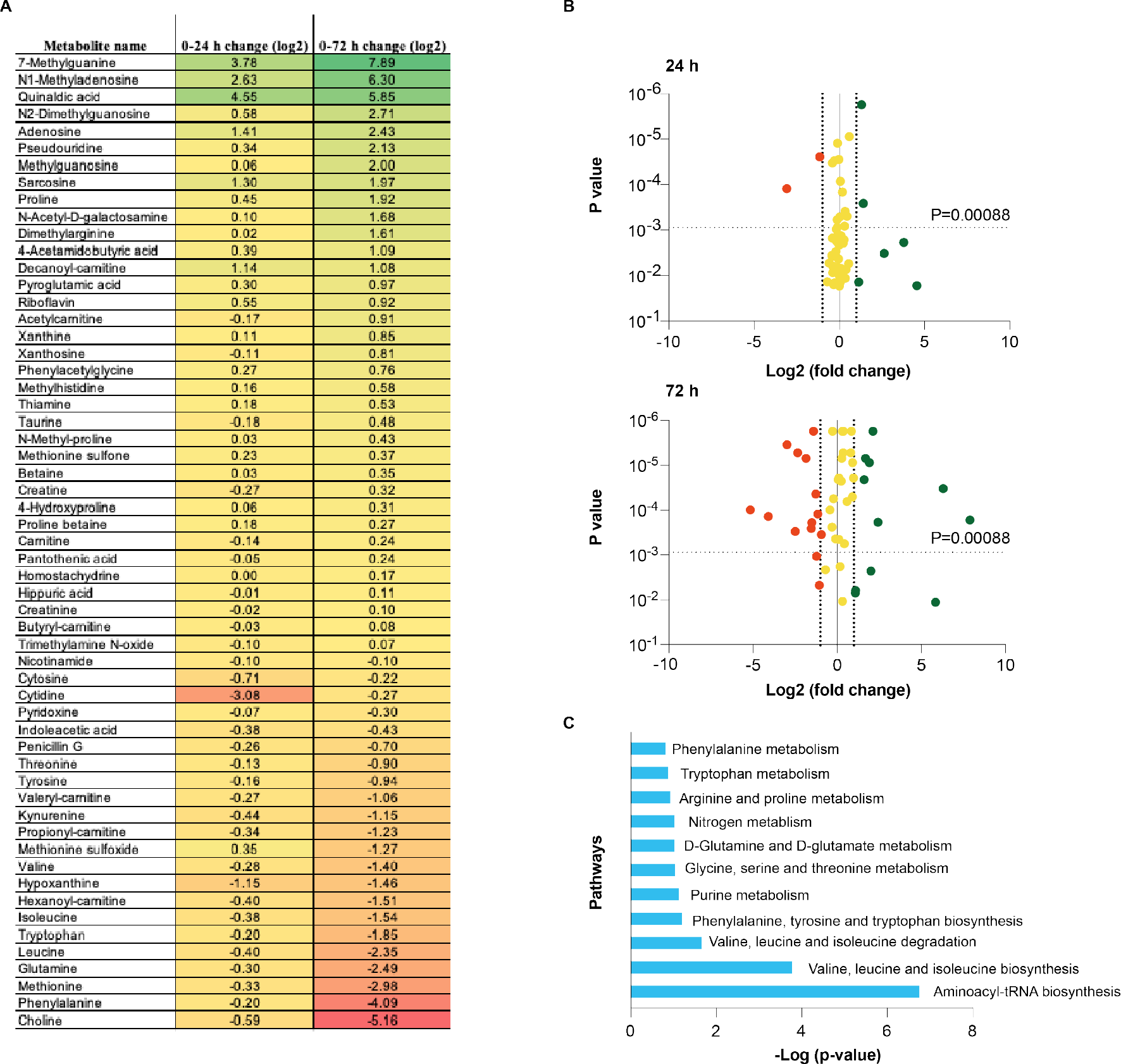
LC-MS-based untargeted metabolic analysis of extracellular metabolites in CCM from HepG2 hepatocytes. One microliter of CCM at 0, 24 and 72 h were extracted using 100 μL of acetonitrile. A, Table showing the folds change (log2 samples: 0 h control) of the 57 annotated metabolites. B, volcano plots of the extracellular metabolites (green: >2-fold increase and red: >2-fold decrease considering p-value <0.00088 (0.05/57)). C, Metaboanalyst pathways analysis of compounds showing >2-fold change and p-value <0.01 at 72 h.

## Acknowledgements

This work was supported by JSPS KAKENHI Grant Number 17H02083, 20K20168 and 19K17662. Kyoto University GAP fund program (207010). We acknowledge the support from the Gunma University Initiative for Advanced Research (GIAR). IM was supported by Japan Society for the Promotion of Science (JSPS) postdoctoral fellowship (P17774). Authors thank Dr. Satoshi Imamura for assisting in HepG2 cells culture.

